# Quorum sensing-induced phenotypic switching as a regulatory nutritional stress response in a competitive two-species biofilm: An individual-based cellular automata model

**DOI:** 10.1101/2020.06.15.153718

**Authors:** Tejesh Reddy Chirathanamettu, Parag D. Pawar

## Abstract

Competition for nutrients in a polymicrobial biofilm may lead to susceptible species being subjected to nutritional stress. The influence of bacterial growth rates and interspecies interactions on their susceptibility and response to nutritional stress is not well understood. *Pseudomonas aeruginosa* and *Staphylococcus aureus* are two prevalent causative pathogens that coexist in biofilm-associated infections. Despite being the slower-growing species, *P. aeruginosa* dominates in a two-species biofilm by inducing phenotypic switching of *S. aureus* to a metabolically-challenged small colony variant (SCV) via the release of 2-heptyl-4-hydroxyquinoline N-oxide (HQNO). We hypothesize that *P. aeruginosa* experiences nutritional stress in competition with *S. aureus*, and that the release of HQNO is an adaptive response to nutritional stress. We present an individual-based two-species biofilm model in which interactions between entities induce emergent properties. As the biofilm matured, the difference in growth rates of the two species caused a non-uniform distribution of nutrients leading to nutritional stress for *P. aeruginosa* and a concurrent increase in the proportion of *S. aureus* subpopulation. The latter resulted in increased release of autoinducer, and subsequently the upregulation of *P. aeruginosa* cells via quorum sensing. Upregulated *P. aeruginosa* cells released HQNO at enhanced rates, thereby inducing phenotypic switching of *S. aureus* to SCVs which consume nutrient at a reduced rate. This shifted the nutrient distribution back in favor of *P. aeruginosa*, thereby relieving nutritional stress. Increase in nutritional stress potentiated the transformation of *S. aureus* into SCVs. HQNO production decreased once nutritional stress was relieved, indicating that phenotypic switching acts as a regulatory stress-adaptive response.

## 1. Introduction

It has been well established that bacteria grow preferentially in the biofilm mode of growth wherein microorganisms form self-assembled, surface-associated communities (Hall-Stoodley *et al*. 2004). Biofilms are associated with recurrent infections that are difficult to treat because of their recalcitrance to antibiotics. *In vitro* studies assessing growth dynamics and antibiotic tolerance of biofilms have typically been performed with single species (Sena *et al*. 2006; Ito *et al*. 2009; Abidi *et al*. 2013; Duus *et al*. 2013; Eyoh *et al*. 2014; Vuotto *et al*. 2014; Moghadam *et al*. 2014; Qi *et al*. 2016). However, bacteria coexist in nature, and under pathological conditions, to form communities that feature advanced synergistic and antagonistic interactions, influencing growth and metabolism of other species (Haruta *et al*. 2009). While monoculture biofilms are usually associated with implant infections (Berbari *et al*. 2006; Gutierrez Jauregui *et al*. 2019), several common infections involve polymicrobial biofilms, including urinary tract (Swidsinski *et al*. 2013; Azevedo *et al*. 2017), respiratory (Høiby *et al*. 2017; McDaniel *et al*. 2020), otitis media (Armbruster *et al*. 2010; Perez *et al*. 2014) and wound infections (Gjødsbøl *et al*. 2006; Gardner and Frantz 2008). In addition, infections involving polymicrobial biofilms have higher mortality rates compared to single-species biofilms (Pulimood *et al*. 2002). Experimental evidence suggests that biofilm-associated bacteria, in response to competitors, alter their behavior by enhancing biofilm formation (Oliveira *et al*. 2015), changing the amount of secreted antimicrobials (Traxler *et al*. 2013; Abrudan *et al*. 2015), or by increasing tolerance towards antibiotics (Parijs and Steenackers 2018). Ecological competition for nutritional resources, especially in polymicrobial biofilms, is often intense (Oliveira *et al*. 2015; Nadell *et al*. 2016), influencing species-wise nutrient availability and uptake, and ultimately the predominance of different taxa (Tilman 1977).

Two bacterial pathogens prevalent in chronic infections that have evolved an intricate system of interactions in their battle for space and nutrients are *Pseudomonas aeruginosa* (*P. aeruginosa*) and *Staphylococcus aureus* (*S. aureus*). Polymicrobial biofilms colonized by these two species are often encountered in airways of patients with cystic fibrosis (CF) (Filkins and O’Toole 2015) and in chronic wounds (Fazli *et al*. 2009). Interactions between the two species are known to be synergistic as well as antagonistic. For instance, co-existence with *S. aureus* protects *P. aeruginosa* by inhibiting its phagocytosis: α toxin inhibits phagocytosis by macrophages in lungs, thereby reducing effective killing of both *S. aureus* and Gram negative bacteria (Cohen *et al*. 2016). *S. aureus* also facilitates survival of *P. aeruginosa* in CF patients by detoxifying surrounding nitric oxide released by host immune cells (Hoffman *et al*. 2010). On the other hand, exoproducts like staphylococcal protein A secreted by *S. aureus* inhibit biofilm formation by *P. aeruginosa* (Armbruster *et al*. 2016). Competition between the two species for iron enhances production of antimicrobials such as alkyl quinolones by *P. aeruginosa* thereby suppressing *S. aureus* growth (Nguyen *et al*. 2015). In addition, cross feeding of metabolic byproducts secreted by other bacteria has been shown to enhance *P. aeruginosa* metabolic competition (Phalak *et al*. 2016).

Recent experimental work investigating the influence of laboratory culture media on *in vitro* biofilm growth suggests that *P. aeruginosa* exhibits lower growth rates compared to *S. aureus*, with the grow rate of co-culture biofilms being greater than that of *P. aeruginosa* monocultures, but less than that of *S. aureus* alone (Wijesinghe *et al*. 2019). Along these lines, the doubling times for *P. aeruginosa* were found to be nearly five times those for *S. aureus* (McBirney *et al*. 2016). The influence of this striking difference in growth rates on the nutritional distribution between the two species in a biofilm setting is not fully understood. We hypothesize that *P. aeruginosa* is at a metabolic disadvantage under these conditions on account of nutritional stress. Stress responses enable individual bacteria to adapt to environmental stresses such as low nutrient availability and treatment with antimicrobial agents. Perhaps partially in response to the metabolic disadvantage, *P. aeruginosa* has evolved a number of mechanisms to enhance its competitiveness in multispecies biofilms. The most widely studied mechanism is growth inhibition and lysis of competing bacteria through secretion of small molecules such as 2-heptyl-4-hydroxyquinoline N-oxide (HQNO) (Mitchell *et al*. 2010) and pyocyanin (Lau *et al*. 2004), and protein toxins including bacteriocins (Bakkal *et al*. 2010).

Quorum sensing (QS) refers to a broad intercellular communication system used by bacteria to collectively coordinate group behavior based on their population density. This process relies on the production, release, and group-wise detection of signal molecules called autoinducers which rapidly diffuse in the aqueous phase and across cell populations, and accumulate in the biofilm over time. In a two-species biofilm, *P. aeruginosa* cells sense an increase in *S. aureus* cell density via the signaling molecule N-acetyl glucosamine (GlcNAc) (Korgaonkar and Whiteley 2011). This Gram-positive cell wall polymer molecule induces Pseudomonas quinolone signal (PQS)-based quorum sensing (Korgaonkar *et al*. 2013). The PQS QS system regulates the pqsABCDE operon which enhances the production of HQNO by *P. aeruginosa* (Déziel *et al*. 2004). Extracellular factors such as HQNO secreted by *P. aeruginosa* have been shown to subjugate *S. aureus* to persist as small colony variants (SCVs) (Hoffman *et al*. 2006). HQNO, which is an anti-staphylococcal compound, has no lytic activity against *S. aureus* itself, but rather slows down its growth by inhibiting oxidative respiration (Hoffman *et al*. 2006) via inhibition of the cytochrome systems (LIGHTBOWN and JACKSON 1956). The effect of HQNO-induced phenotypic switching of wild-type *S. aureus* to SCVs on biofilm physiology is not fully understood.

In addition to biochemical factors, interspecies interactions may also be influenced by the physical architecture of biofilms. Biofilms are spatially heterogeneous with cell behavior being tightly regulated by the local fluid environment, reaction-diffusion barriers, and interactions with neighboring cells. Concentration gradients due to diffusion-limitation establish heterogeneous niches within the biofilm that can produce spatial variation in the biomass density in monoculture (Stewart 1998; Machineni *et al*. 2017, 2018) and multi-species biofilms (Woods *et al*. 2012; Mazumdar *et al*. 2013). Several computational studies have modeled the growth dynamics of monoculture biofilms to investigate the influence of spatial heterogeneity (Machineni *et al*. 2017), cell death (Fozard *et al*. 2012), dispersal (Emerenini *et al*. 2015), and QS (Langebrake *et al*. 2014). Recently, mathematical models have been developed for multi-species biofilms studying the behavior of the residing species and their interactions over available nutrients (Phalak *et al*. 2016; Martin *et al*. 2017). A 1D metabolic model was developed for a two-species biofilm system predicting spatial partitioning among the two species, and lysis of one species by secretions of the other (Phalak *et al*. 2016); the model uses a fixed biofilm thickness, and does not consider the stochastic nature of biofilm growth. A 2D cellular automata model for a two-species oral biofilm was used to investigate the competition for nutrients, but only during initial growth phases (Martin *et al*. 2017). In the past, influence of interspecies interactions via QS and phenotypic switching, and their influence on nutritional stress have not been considered.

Development of a polymicrobial biofilm involves interplay between complex, stochastic processes including interspecies interactions such as QS and phenotypic switching. Furthermore, biofilms are spatially heterogeneous, with their growth dynamics strongly influenced by growth rates of individual bacterial cells and the local nutrient availability. Local nutrient concentrations, in turn, depend on the nutrient concentration in the bulk phase, and on biochemical properties which influence nutrient diffusion in the biofilm (Stewart 2003; Petroff *et al*. 2011; Guélon *et al*. 2012). In this work, we present a 3D individual-based cellular automata approach to simulate the growth dynamics of a two-species bacterial biofilm comprising the wild-type species *P. aeruginosa* and *S. aureus*. Such individual-based models represent bacterial cells as discrete units, allowing for variability in properties and behavior, including growth rates, nutrient uptake, upregulation and downregulation states, QS signal secretion, HQNO production, and stochastic replication according to a set of predetermined rules. Key cellular processes incorporated in the model include nutrient transport and uptake, bacterial growth, division, death, detachment, interspecies QS, and phenotypic switching are incorporated in the model. The discrete and 3D individual-based nature of the model, combined with physical dynamics, cause heterogeneities in nutrient availability leading to development of nutritional stress as a consequence of interactions with the local environment and between the species, rather than being a model input.

How bacteria respond to the presence of competing strains in mixed-species biofilms under varying nutrient conditions is not fully understood. In this work, we investigate influence of competition for nutritional resources between coexisting bacterial strains on the growth dynamics of a mixed-species biofilm. We hypothesize that the slower-growing *P. aeruginosa* uses HQNO-induced transformation of wild-type *S. aureus* to SCVs as a mechanism of defense against nutritional stress. Our results indicate that *P. aeruginosa*, the slower growing of the two species, is under nutritional stress in competition with *S. aureus*. This triggers the release of HQNO via a QS-dependent mechanism, and the subsequent transformation of wild-type *S. aureus* into SCVs which exhibit a lower nutrient uptake rate. This, in turn, relieves the nutritional stress for *P. aeruginosa*. We show that conditions involving higher nutritional stress for *P. aeruginosa* result in enhanced transformation of wild-type *S. aureus* into SCVs. The results also illustrate that this mechanism is reversible: that is, once nutritional stress for *P. aeruginosa* is relieved, HQNO production slows down. To our knowledge, this is the first individual-based biofilm computational framework that incorporates phenotypic switching from wild-type cells into SCVs. Understanding the spatio-temporal dynamics of interspecies interactions in polymicrobial biofilms may provide clues into the underlying mechanisms of enhanced antibiotic resistance.

## 2. Methods

### 2.1 Domain geometry and model components

We have previously described the 3D rectangular domain used in this model (Machineni *et al*. 2017, 2018). The bottom plane is a square with side 120 µm and represents the stationary substratum upon which the biofilm develops. A continuously replenished nutrient reservoir is placed at the top at a constant distance from the substratum. The interface between the reservoir and the biofilm domain is termed the diffusion boundary layer (DBL). The DBL is assumed to remain parallel to the substratum, and has a constant thickness of 18 µm. The space between the DBL and the substratum is discretized into cubical elements of volume 27 µm^3^ each. Each element may be occupied by one or more of the following entities: (i) bacterial cells (wild-type *S. aureus*, wild-type *P. aeruginosa*, and small colony variant of *S. aureus*) and (ii) dissolved entities (nutrient, HQNO, and autoinducer). Whereas each cubical element can be occupied by only one bacterial cell, dissolved entities are capable of coexisting with each other and with the bacteria cell in the same cubical element. Periodic boundary conditions are employed in the horizontal directions, thereby eliminating edge effects and ensuring continuity of biomass (Chang *et al*. 2003; Picioreanu *et al*. 2004).

Each bacterial cell in the domain is modeled as an independent unit with its own set of parameters (Table 1) and behaviors. To simulate variability between individual cells, parameter values were obtained by random draws from a uniform distribution around the average values listed in Table 1. Negative values and those outside ±10% of the mean were discarded (Kreft *et al*. 1998). Investigations involving individual-based biofilm models have shown that simulations involving up to 10,000 bacteria are sufficient to reproduce all steps of biofilm formation observed in experiments. Using our domain geometry we were able to simulate biofilms comprising of up to 24711 ± 68 bacterial cells of all species taken together. The resulting aggregate behavior of the biofilm is emergent from the local interactions between the individual bacteria, and their surroundings, thereby allowing us to simulate the self-organized process of biofilm formation.

**Table1:**
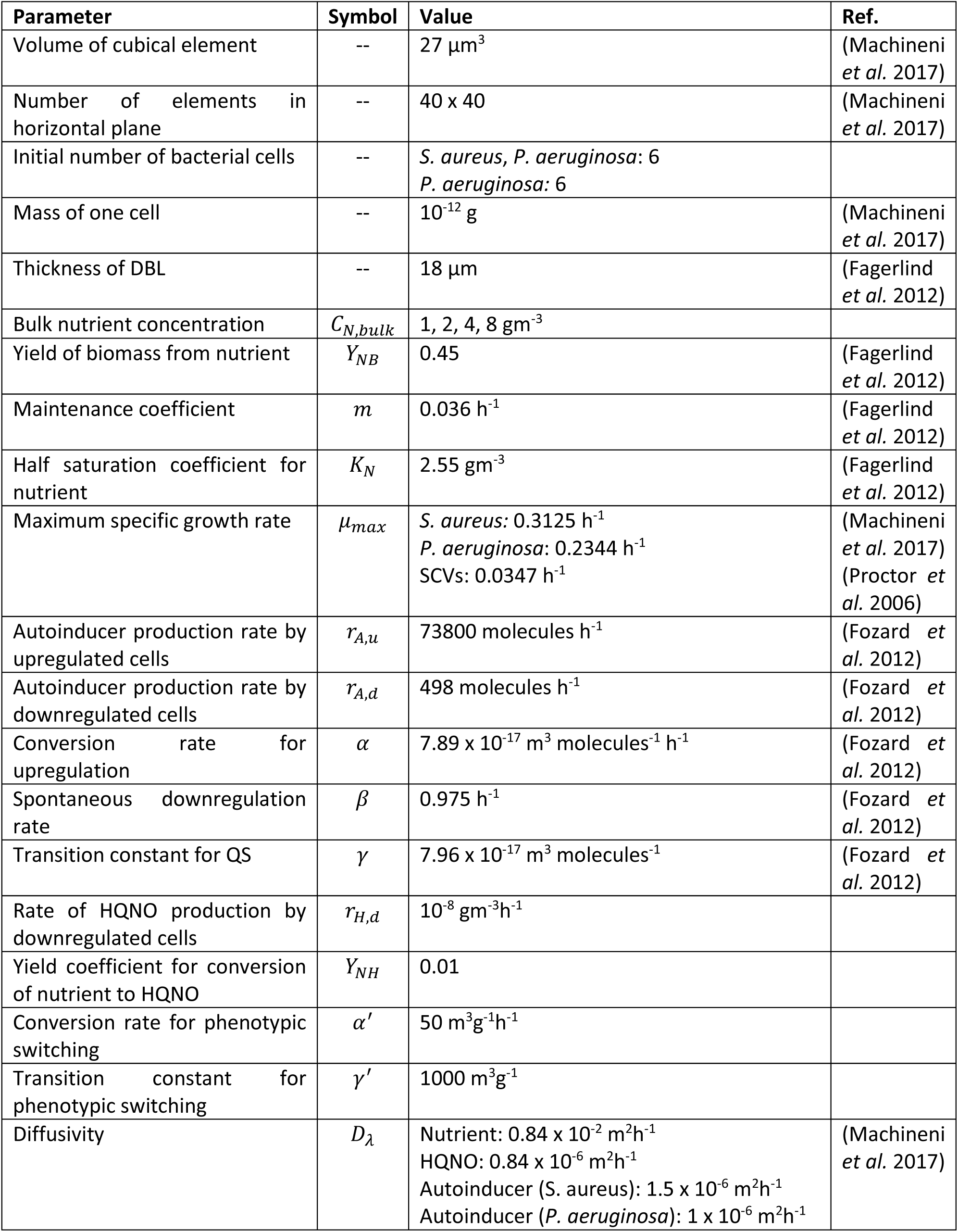
Model parameters.

| Parameter | Symbol | Value | Ref. |
| --- | --- | --- | --- |
| Volume of cubical element | -- | $27 \mu\text{m}^3$ | (Machineni *et al.* 2017) |
| Number of elements in horizontal plane | -- | 40 x 40 | (Machineni *et al.* 2017) |
| Initial number of bacterial cells | -- | *S. aureus*, *P. aeruginosa*: 6<br>*P. aeruginosa*: 6 |  |
| Mass of one cell | -- | $10^{-12} \text{ g}$ | (Machineni *et al.* 2017) |
| Thickness of DBL | -- | $18 \mu\text{m}$ | (Fagerlind *et al.* 2012) |
| Bulk nutrient concentration | $C_{N,bulk}$ | 1, 2, 4, 8 $\text{gm}^{-3}$ | |
| Yield of biomass from nutrient | $Y_{NB}$ | 0.45 | (Fagerlind *et al.* 2012) |
| Maintenance coefficient | $m$ | $0.036 \text{ h}^{-1}$ | (Fagerlind *et al.* 2012) |
| Half saturation coefficient for nutrient | $K_N$ | $2.55 \text{ gm}^{-3}$ | (Fagerlind *et al.* 2012) |
| Maximum specific growth rate | $\mu_{max}$ | *S. aureus*: $0.3125 \text{ h}^{-1}$<br>*P. aeruginosa*: $0.2344 \text{ h}^{-1}$<br>SCVs: $0.0347 \text{ h}^{-1}$ | (Machineni *et al.* 2017)<br>(Proctor *et al.* 2006) |
| Autoinducer production rate by upregulated cells | $r_{A,u}$ | $73800 \text{ molecules h}^{-1}$ | (Fozard *et al.* 2012) |
| Autoinducer production rate by downregulated cells | $r_{A,d}$ | $498 \text{ molecules h}^{-1}$ | (Fozard *et al.* 2012) |
| Conversion rate for upregulation | $\alpha$ | $7.89 \times 10^{-17} \text{ m}^3 \text{ molecules}^{-1} \text{ h}^{-1}$ | (Fozard *et al.* 2012) |
| Spontaneous downregulation rate | $\beta$ | $0.975 \text{ h}^{-1}$ | (Fozard *et al.* 2012) |
| Transition constant for QS | $\gamma$ | $7.96 \times 10^{-17} \text{ m}^3 \text{ molecules}^{-1}$ | (Fozard *et al.* 2012) |
| Rate of HQNO production by downregulated cells | $r_{H,d}$ | $10^{-8} \text{ gm}^{-3}\text{h}^{-1}$ | |
| Yield coefficient for conversion of nutrient to HQNO | $Y_{NH}$ | 0.01 | |
| Conversion rate for phenotypic switching | $\alpha'$ | $50 \text{ m}^3\text{g}^{-1}\text{h}^{-1}$ | |
| Transition constant for phenotypic switching | $\gamma'$ | $1000 \text{ m}^3\text{g}^{-1}$ | |
| Diffusivity | $D_\lambda$ | Nutrient: $0.84 \times 10^{-2} \text{ m}^2\text{h}^{-1}$<br>HQNO: $0.84 \times 10^{-6} \text{ m}^2\text{h}^{-1}$<br>Autoinducer (*S. aureus*): $1.5 \times 10^{-6} \text{ m}^2\text{h}^{-1}$<br>Autoinducer (*P. aeruginosa*): $1 \times 10^{-6} \text{ m}^2\text{h}^{-1}$ | (Machineni *et al.* 2017) |

The simulation represents a time march in which the occupancy state of each element is updated at every time step. Initially, six cells each of wild-type *S. aureus* and wild-type *P. aeruginosa* are placed into random elements on the substratum. Nutrient diffuses across the DBL, and is consumed by the cells. Subsequently cells grow, divide, die, and detach, resulting in the formation of a contiguous multicellular population that comprises the two-species biofilm. At the end of each time step, the nutrient reservoir is shifted vertically upwards so as to maintain the predetermined thickness of the DBL. In select runs, bacterial cells of the two wild-type species were allowed to secrete autoinducer molecules and exhibit QS behavior. In such runs, *P. aeruginosa* cells released HQNO, which induced phenotypic switching resulting in the transformation of wild-type *S. aureus* cells into SCVs.

### 2.2 Reaction and Diffusion

The model incorporates the following dissolved entities: (i) nutrient, (ii) autoinducer, and (3) HQNO. These are either consumed (nutrient) or secreted (autoinducer, HQNO) by the bacterial cell residing in the same cubical element, resulting in the generation of local concentration gradients. In turn, the spatio-temporal variation of these concentrations influences biomass growth as well as cellular behavior and inter-cellular interactions. The concentration of each dissolved entity within each element of the domain changes because of diffusion and uptake or release by bacterial cells. The concentration field *C*_*λ*_ = *C*_*λ*_(*x*_1_, *x*_2_, *x*_3_, *t*) of dissolved entity *λ* is governed by the reaction-diffusion equation

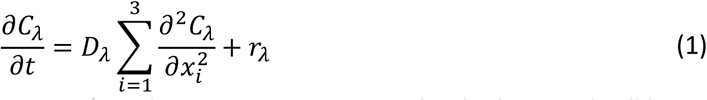

in which *D*_*λ*_ is the diffusivity, and *r*_*λ*_ is the rate of production or consumption by the bacterial cell located at location (*x*_1_, *x*_2_, *x*_3_). The 3D reaction-diffusion equation is solved numerically with the following boundary conditions—

a. A Dirichlet boundary condition is imposed at the DBL, i.e., the concentration remains constant at the interface between the boundary layer and bulk liquid. This constant bulk concentration is predetermined for the nutrient (C_N,bulk_), and is set to zero for autoinducer and HQNO.
b. Neumann boundary condition is imposed at the substratum, where the flux is zero.
c. Periodic boundary conditions are applied at the lateral boundaries.

### 2.3 Nutrient uptake

The nutrient (*λ* = *N*) uptake rate (*r*_*N*_) is a function of the bacterial biomass *C*_*B*_(*x*_1_, *x*_2_, *x*_3_, *t*) and nutrient concentration *C*_*B*_(*x*_1_, *x*_2_, *x*_3_, *t*), and is described by the Herbert-Pirt model.

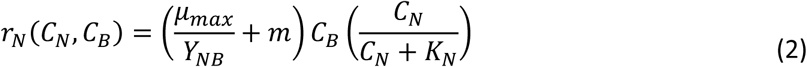

Here, *μ*_*max*_, *Y*_*NB*_, and *m* respectively represent the maximum specific growth rate, yield coefficient, and maintenance coefficient of bacteria, and *K*_*N*_ is the half-saturation coefficient of the nutrient.

### 2.4 Cellular processes

Modeling of cellular behavior including growth, division, death, and detachment has been described in detail elsewhere (Machineni *et al*. 2017, 2018). Here, we present a brief description of these processes along with the governing equations.

#### 2.4.1 Growth

A portion of the consumed nutrient is utilized by the bacterium towards endogenous metabolism; this is assumed to be proportional to the biomass concentration. Leftover nutrient is converted to biomass with an efficiency represented by the yield coefficient, *Y*_*NB*_. The net accumulation of biomass is given by

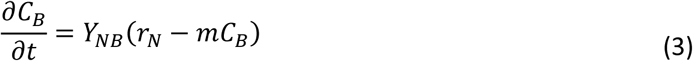

#### 2.4.2 Division

A bacterial cell divides into two daughter cells when its biomass reaches twice its native value. Whereas one daughter cell continues to occupy the same element as the mother cell, the other is inserted into a bacterium-free element in the immediate Moore neighborhood. If multiple bacterium-free elements are available for occupation, one is chosen at random (Picioreanu *et al*. 1998). If all elements in the Moore neighborhood are occupied by bacteria, a nearest unoccupied element is identified based on its Chebyshev distance from the mother cell, and bacterial cells that lie between the mother cell and this unoccupied element are shifted towards the empty element, thereby creating a bacterium-free element in the Moore neighborhood of the mother cell. This newly created bacterium-free element is then occupied by the daughter cell.

#### 2.4.3 Death

Cell death is assumed to occur via one of two mechanisms: (i) limited nutrient uptake (Hunt *et al*. 2004) or (ii) starvation caused by prolonged stay in the stationary phase (Nyström 2001, 2003). Nutrient uptake is quantified by the ratio (*R*) defined as the ratio of rate of nutrient uptake to that of endogeneous metabolism.

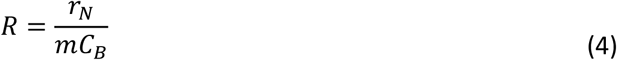

Cell death by limited nutrient uptake is assumed to occur when *R* falls below a certain threshold (*R*_*min*_ = 0.15). If *R* falls below 1, the cell exhibits zero or negative net growth, and is said to have entered the stationary phase. Cell death is assumed to occur if the cell continues to remain in this growth-arrested phase for a preset number of hours (*t*_*SP*_ = 24 *h*). Dead cells are discarded from the simulation domain and are no longer tracked.

#### 2.4.4 Detachment

Cell detachment is modeled using a simplified geometrical model governed by physical connectivity of cells to the substratum. Bacteria are connected to the substratum either directly or indirectly through a group of live bacteria in which at least one bacterium is directly bound to the substratum. Connectivity of cells to the substratum is influenced by localized cell death events. At the end of each time step, detachment events are recorded, and detached cells are removed from the domain.

### 2.5 Quorum sensing and secretion of autoinducer

In select runs, wild-type bacterial cells secrete autoinducer, and exhibit QS behavior by altering their states dependent on local autoinducer concentration. Bacteria are modeled as being in either upregulated or downregulated states, with the latter being the default. Cells switch between these states at rates dependent on the local autoinducer concentration, *C*_*A*_(*x*_1_, *x*_2_, *x*_3_, *t*). The probabilities of switching from one state to another within a time interval Δ*t* are calculated based on the respective transition rates.

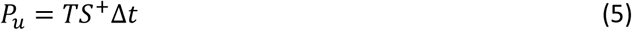

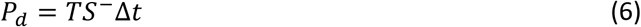

Here, *P*_*u*_ and *P*_*d*_ represent the probability of upregulation and downregulation, respectively. *TS*^+^ and *TS*^−^ are the transition rates of transformation from downregulated to upregulated and from upregulated to downregulated states, respectively. The transition rates, in turn, are functions of the autoinducer concentration, and are described by Holling’s Type II functional response.

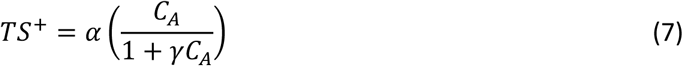

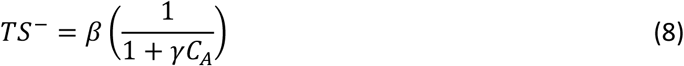

where *α* and *β* are the spontaneous upregulation and downregulation rates, and *γ* is the transition constant. To determine whether a bacterium switches between states, the probability of switching is compared with a random number (*n*_*R*_) generated from a uniform distribution on the interval [0,1]. The bacterium switches states if the probability of switching is greater than *n*_*R*_.

Upregulated and downregulated cells are assumed to secrete autoinducer molecules (*λ* = *A*) at constant rates written as

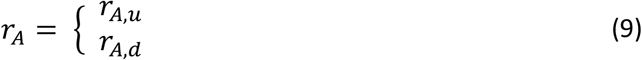

in which *r*_*A,u*_ > *r*_*A,d*_.

### 2.6 Phenotypic switching and HQNO release

In biofilms that exhibit QS, *P. aeruginosa* cells secrete HQNO which induces wild-type *S. aureus* cells to undergo phenotypic switching to the SCV phenotype dependent on the local HQNO concentration. Such biofilms are henceforth referred to as SCV^+^. The probability of phenotypic switching (*P*_*SCV*_) within a time interval Δ*t* is calculated based on a transition rate of switching.

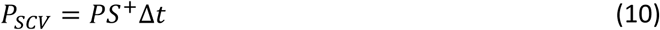

in which *PS*^+^represents the transition rate of phenotypic switching from a single wild-type *S. aureus* cell to a SCV. The transition rate is a function of the local HQNO concentration (*C*_*H*_).

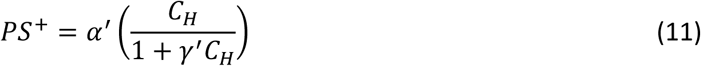

where *α*^′^ and *γ*^′^ are the spontaneous rate of switching and transition constant, respectively. Phenotypic switching occurs if *P*_*SCV*_ > *n*_*R*_, where *n*_*R*_ is a random number generated from a uniform distribution on the interval [0,1]. Whereas downregulated *P. aeruginosa* cells produce HQNO (*λ* = *H*) at a slow, constitutive rate (*r*_*H,d*_), upregulated *P. aeruginosa* cells produce HQNO at an enhanced rate, *r*_*H,u*_, given by

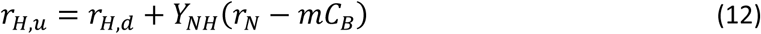

in which *Y*_*NH*_ represents the yield coefficient for conversion of unutilized nutrient into HQNO.

### 2.7 Model simulation and numerical scheme

The simulation represents a time march in which the occupancy state of each cubical element is updated at discrete time steps of 1 h. The model processes described above can be broadly classified into two categories based on their timescales. A multiscale integration approach with two distinct timescales is, therefore, employed: (1) an outer loop with a time step of 1 h in which “slower” cellular processes such as biomass growth, division, death, detachment, and switching between states are tracked, and (2) an inner loop with a finer resolution of 1 x 10^−6^ h where the reaction-diffusion equations are solved to determine equilibrium concentrations of dissolved entities (Eq. 1). Numerical solutions to the reaction-diffusion equations are obtained using a second-order forward-time central-space scheme. The Java programming language is used since it provides a convenient object-oriented framework well-suited for the individual-based model described here.

## 3. Results and Discussion

### 3.1 Biofilm growth dynamics of a two-species biofilm: influence of nutrient concentration

Previously, we have used our model to investigate the roles of nutrient availability and QS on the growth dynamics of a monomicrobial biofilm (Machineni *et al*. 2017, 2018). In the present work, we extend the model to a two-species biofilm comprising of *P. aeruginosa* and *S. aureus*, competing for nutrients. Interspecies interactions are simulated using an autoinducer-based QS mechanism. In select runs, wild-type *S. aureus* cells were allowed to transform into a metabolically challenged phenotype, viz. SCVs, dependent on the local HQNO concentration. For brevity, such biofilms are referred to as SCV^+^, whereas biofilms in which phenotypic switching doesn’t occur are referred to as SCV^-^. To simulate varied nutrient conditions, biofilms were subjected to a range of bulk nutrient concentrations ranging from 1 gm^-3^ to 8 gm^-3^ (Fagerlind *et al*. 2012).

As a first step, we simulated the growth dynamics of a SCV^-^ biofilm; this is analogous to experimental two-species biofilm systems in which a HQNO^-^ mutant strain of *P. aeruginosa* is used, and serves as control for SCV^+^ biofilms (Filkins *et al*. 2015). To delineate the influence of nutrient availability on growth dynamics, we tracked the total accumulated biomass for biofilms subjected to a range of bulk nutrient concentrations. In close agreement with experimental observations (Bester *et al*. 2005; Kroukamp *et al*. 2010), the model was able to simulate five distinct growth phases: lag phase, exponential growth, stationary phase, sloughing, and regrowth (Kroukamp *et al*. 2010) (Fig. 1a). Biofilm growth exhibited a short lag phase whose duration related inversely with the bulk nutrient concentration (20 h, 25 h, 35 h, and 60 h for C_N,bulk_ = 8 gm^-3^, 4 gm^-3^, 2 gm^-3^, and 1 gm^-3^, respectively) during which cell numbers were low (less than 150). This was followed by the exponential phase in which growth rates and peak biomass were higher for biofilms associated with higher bulk nutrient concentrations; for instance, the growth rate for C_N,bulk_ = 8 gm^-3^ was 3.9 x 10^−4^ µg/h, whereas that for C_N,bulk_ = 1 gm^-3^ was 5.4 x 10^−5^ µg/h. Increase in cell population results in increased nutrient consumption, causing the average nutrient concentration within the biofilm domain to drop rapidly (Fig. 1b). We propose that this decrease in nutrient concentration during the exponential phase is a key reason for the emergence of nutritional stress in the biofilm.

**Figure 1.**
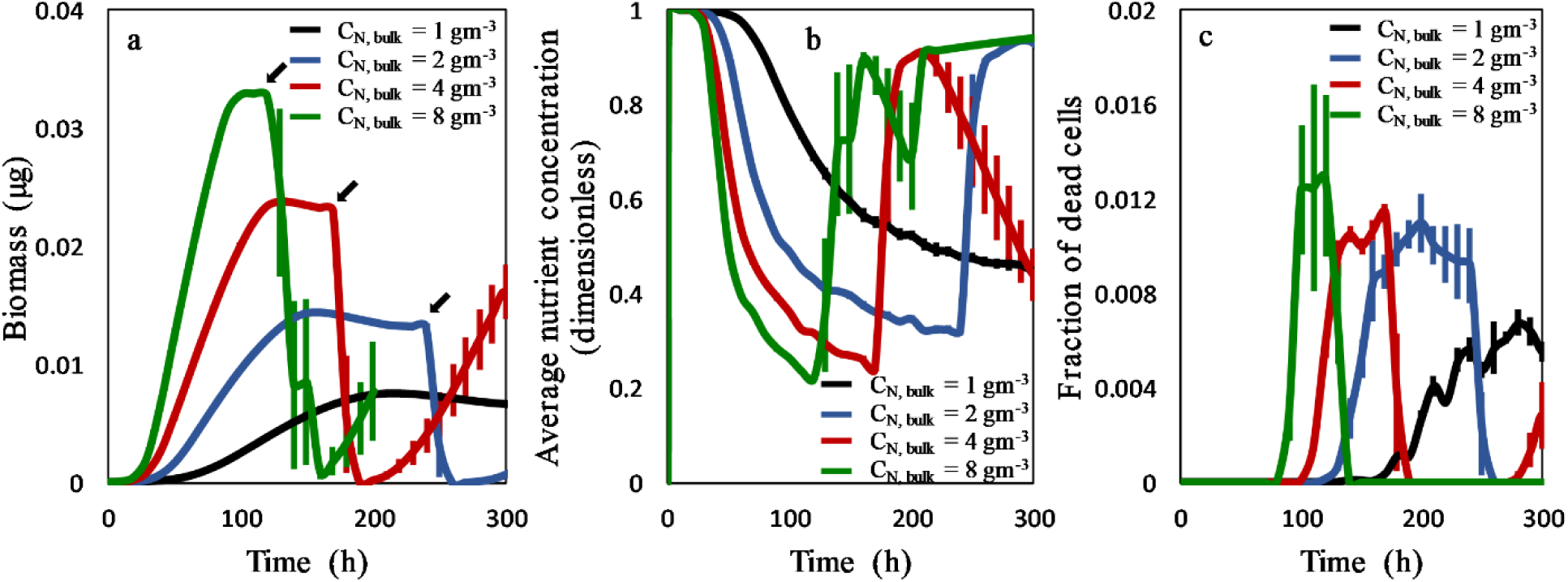
Growth dynamics of a two-species biofilm. Comparison of the development of total biomass (a), average nutrient concentration (b), and fraction of dead cells (c) for bulk nutrient concentrations of 8 gm^-3^ (green), 4 gm^-3^ (red), 2 gm^-3^ (blue), and 1 gm^-3^ (black). The arrows in panel (a) mark the end of the stationary phase and the initiation of sloughing. Data represent mean ± standard error of mean (SEM) of four separate simulations.

Exponential growth subsequently led to the stationary phase in which a dynamic equilibrium was attained between rates of cell division and death, whereupon net biomass remained virtually unchanged. Highest fractions of dead cells for all biofilms were observed in the stationary phase (Fig. 1c). Ultimately, cell death led to detachment events, culminating in the sloughing of the entire biofilm. Interestingly, the lifetime of the fastest growing biofilm (C_N,bulk_ = 8 gm^-3^) was lower (sloughing occurred at ∼140 h) compared to that of the slower growing ones (sloughing for C_N,bulk_ = 4 gm^-3^ occurred at ∼170 h, and that at C_N,bulk_ = 2 gm^-3^ occurred at ∼270 h). In close agreement with our previous observations of the growth dynamics of monomicrobial biofilms (Machineni *et al*. 2017), the two-species biofilm associated with C_N,bulk_ = 1 gm^-3^ exhibited a prolonged stationary phase in which both the total biomass (Fig. 1a) and the fraction of dead cells (Fig. 1c) remained virtually constant, indicating that a balance was established between the rates of biomass formation and depletion. Sloughing did not occur under these conditions. This could be a consequence of the fact that under these nutrient limited conditions, the biofilm thickness is low (data not shown), allowing nutrient to penetrate to the lowest layers adjacent to the substratum. This, in turn, ensures the presence of live cells at the bottom of the biofilm to keep it attached to the substratum at all times. In all cases with sloughing, regrowth of the biofilm occurred post-sloughing. There is considerable variability between results across simulation runs in the regrowth phase due to the high sensitivity to the conditions post-sloughing, i.e. the number of cells that survive detachment.

Fig. 2 shows a representative time evolution of a SCV^+^ biofilm associated with a bulk nutrient concentration of 4 gm^-3^, illustrating the formation of a distinct 3D macrostructure as the biofilm matured, and the various growth stages including: (i) colonization, (ii) early exponential phase, (iii) late exponential phase, (iv) maturation, (v) sloughing, and (vi) regrowth. Early growth phases were associated with distinct colonies of bacteria comprising of primarily only one of the two wild-type species. As the biofilm matured, smaller colonies merged into larger mixed conglomerates in which both species coexisted. The emergence of SCVs was seen during the early exponential growth phase in which cell population is high enough to trigger QS. The representative 2D cross-sections illustrate the spatial distribution of the three species in the biofilm, indicating that whereas wild-type *S. aureus* preferentially localized near the distal regions of the biofilm, *P. aeruginosa* was localized closer to the substratum.

**Figure 2.**
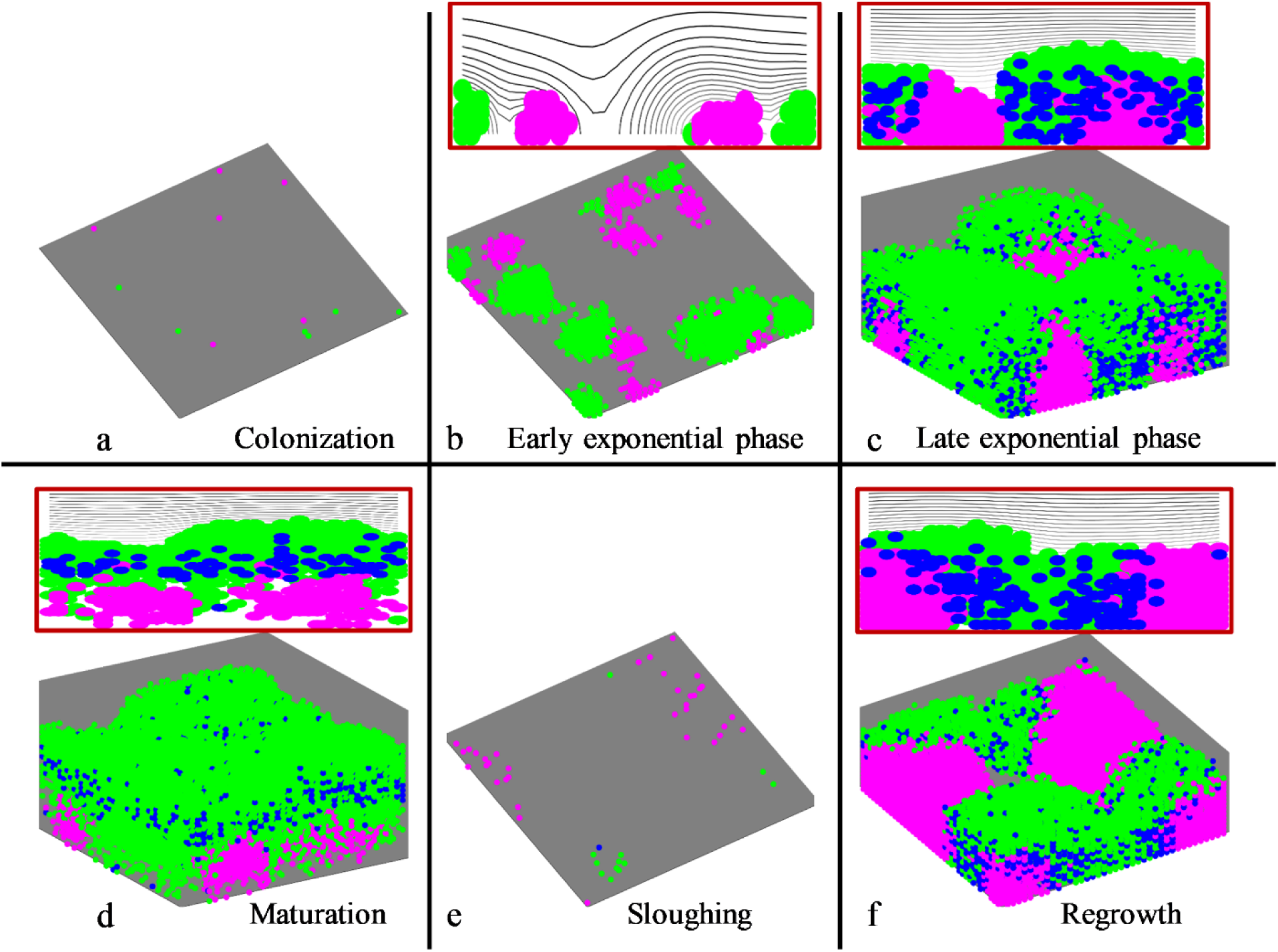
Growth dynamics of a SCV^+^ biofilm. Representative 3D renderings of the time evolution of a SCV^+^ biofilm for C_N,bulk_ = 8 gm^-3^, illustrating different phases of growth at 0 h (a), 30 h (b), 77 h (c), 119 h (d), 139 h (e), and 219 h (f). The biofilm comprises of three species: (i) wild-type *P. aeruginosa* (pink), (ii) wild-type *S aureus* (green), and the SCV phenotype (blue). The insets in panels (b), (c), (d), and (f) show representative 2D cross-sections of the corresponding biofilm to illustrate spatial distribution of the three species.

### 3.2 Interspecies competition induces nutritional stress in P. aeruginosa

Polymicrobial biofilms are associated with intense ecological competition amongst bacterial strains, especially for nutritional resources. We wished to investigate the impact of difference in growth rates of *P. aeruginosa* and *S. aureus* on the relative dominance of the two species under varied nutrient conditions. To this end, we tracked species-wise biomass and the distribution of nutrient uptake between the two species in SCV^-^ biofilms. Species-wise biomass is reported as the relative *P aeruginosa* biomass normalized with the *S. aureus* biomass under the same conditions; this is a simple quantitative measure that correlates inversely with the relative dominance of the latter species in the biofilm. Initially, this biomass ratio is equal to one as both species start with the same number of initial colonizers (6 cells each). As the biofilm matured, the ratio decreased monotonically with time under all bulk nutrient conditions due to the faster growth rates associated with *S. aureus* cells (Fig. 3a). The decrease in the relative biomass of *P. aeruginosa* happens rapidly (for instance, after ∼40 h of growth, the normalized *P. aeruginosa* biomass drops to ∼0.38 for C_N,bulk_ = 8 gm^-3^), and becomes more pronounced as the biofilm matures (after ∼120 h of growth, the biomass ratio drops to less than 0.1 for C_N,bulk_ = 8 gm^-3^). The dominance of *S. aureus* was more pronounced at higher bulk nutrient concentrations. In addition to the higher growth rates associated with *S. aureus*, this marked dominance could also be attributed to preferential localization near the top layers of the biofilm where nutrient availability is higher; in contrast, *P. aeruginosa* cells are localized in close proximity to the substratum with limited access to nutrient (data not shown).

**Figure 3.**
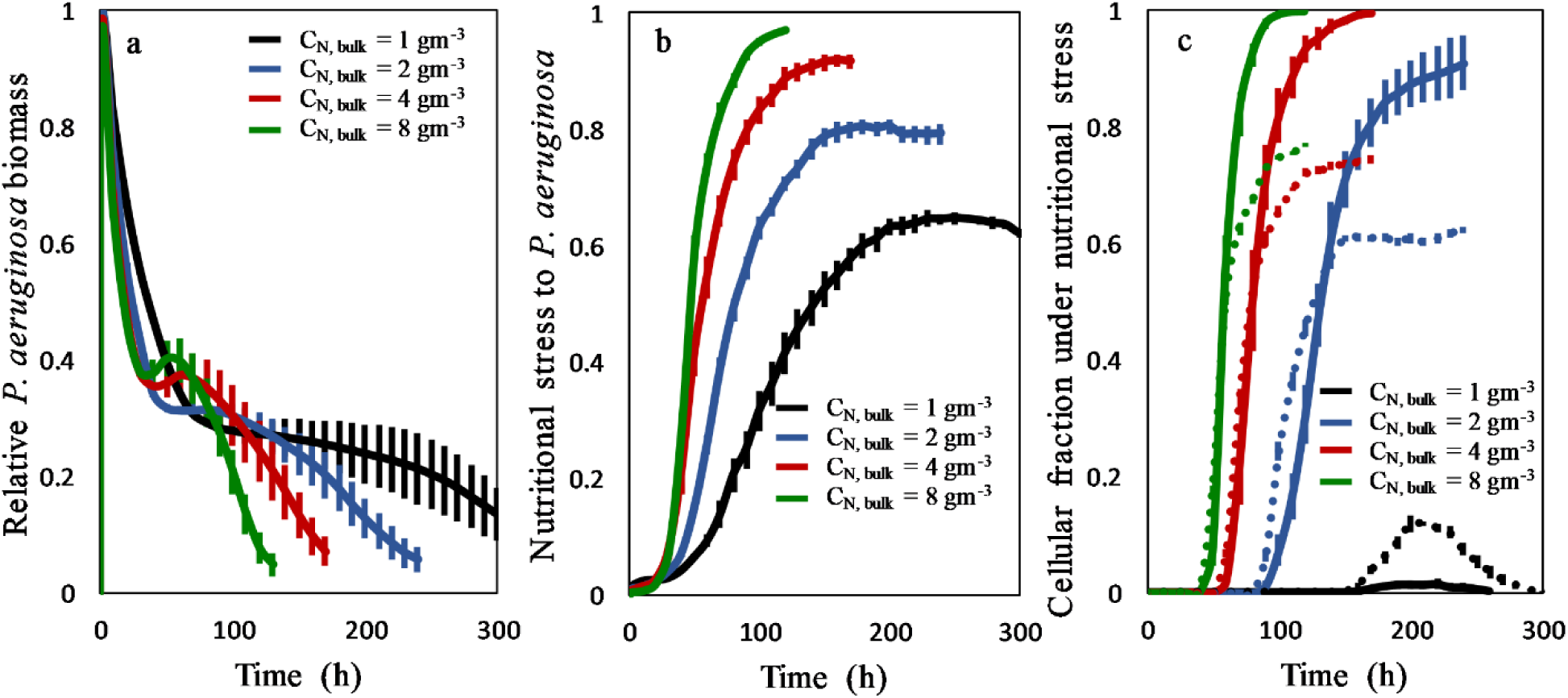
Interspecies competition induces nutritional stress in *P. aeruginosa*. Time evolution of the relative *P. aeruginosa* biomass (a), nutritional stress for *P. aeruginosa* cells (calculated as the difference in the average nutrient availability for all cells in the biofilm minus the nutrient availability for *P. aeruginosa* cells, normalized with the average nutrient availability) (b), and species-wise fraction of cells for which local nutrient concentration is less than 10% of the highest bulk nutrient concentration of 8 gm^-3^ (c) for bulk nutrient concentrations of 8 gm^-3^ (green), 4 gm^-3^ (red), 2 gm^-3^ (blue), and 1 gm^-3^ (black). The solid and dotted lines in (c) represent *P. aeruginosa* and *S. aureus* respectively. Data represent mean ± SEM of four separate simulations.

Next, we quantified nutritional stress associated with *P. aeruginosa* cells as the difference between the average nutrient consumption by all cells and that consumed only by *P. aeruginosa* cells, normalized with the average nutrient consumption by all cells. A value of zero corresponds to *P. aeruginosa* cells being under no nutritional stress (i.e., nutrient is shared equally between the two species). On the other hand, a value of one indicates maximal nutritional stress for *P. aeruginosa* where all nutrient is consumed by *S. aureus* cells in the biofilm. Nutritional stress for *P. aeruginosa* cells was positive under all conditions tested, and increased monotonically with time (Fig. 3b). This is in line with the previous observation that *S. aureus* is the more dominant species in the biofilm (Fig. 3a). Interestingly, *P. aeruginosa* cells in biofilms subjected to higher bulk nutrient concentrations were under greater nutritional stress. This is in agreement with the observation that relative *P. aeruginosa* biomass is inversely related to the bulk nutrient concentration (Fig. 3a). Along these lines, the fraction of *P. aeruginosa* cells with nutrient availability less than 10% of the highest bulk nutrient concentration (C_N,bulk_ = 8 gm^-3^) was higher than that of *S. aureus* cells (Fig. 3c).

### 3.3. Nutritional stress promotes quorum sensing and HQNO production in P. aeruginosa

We hypothesized that *P. aeruginosa*, in an adaptive response to nutritional stress, induces transformation of wild-type *S. aureus* to the slower-growing SCV phenotype. As a first step towards testing this hypothesis, we simulated SCV^+^ biofilms in which the two species were allowed to interact via QS; in such biofilms, *P. aeruginosa* cells secrete HQNO which induces transformation of wild-type *S. aureus* to the slower-growing SCV phenotype. Cells of both wild-type species produce and locally release autoinducer molecules which subsequently spread throughout the biofilm via diffusion. As the biofilm matures, autoinducer concentration in the biofilm increases with increasing cell numbers. In agreement with experimental observations, after the autoinducer concentration reached a threshold value, nearby bacterial cells transitioned to the upregulated state resulting in enhanced production and release of autoinducer molecules. In addition, upregulated *P. aeruginosa* cells also produced HQNO at a higher rate compared to that produced by downregulated *P. aeruginosa* cells. In accordance with experimental observations, the fraction of upregulated cells increased during the exponential growth phase until virtually the entire biofilm rapidly switched from low to high QS activity (Fig. 4a). This switch was delayed under low nutrient supply conditions (C_N,bulk_ = 1 gm^-3^). The average autoinducer concentration increased monotonically with time for all bulk nutrient concentrations (Fig. 4b). Upregulated *P. aeruginosa* cells secreted HQNO at an enhanced rate. Consequently, the average HQNO concentration increased with time for all bulk nutrient concentrations (Fig. 4c).

**Figure 4.**
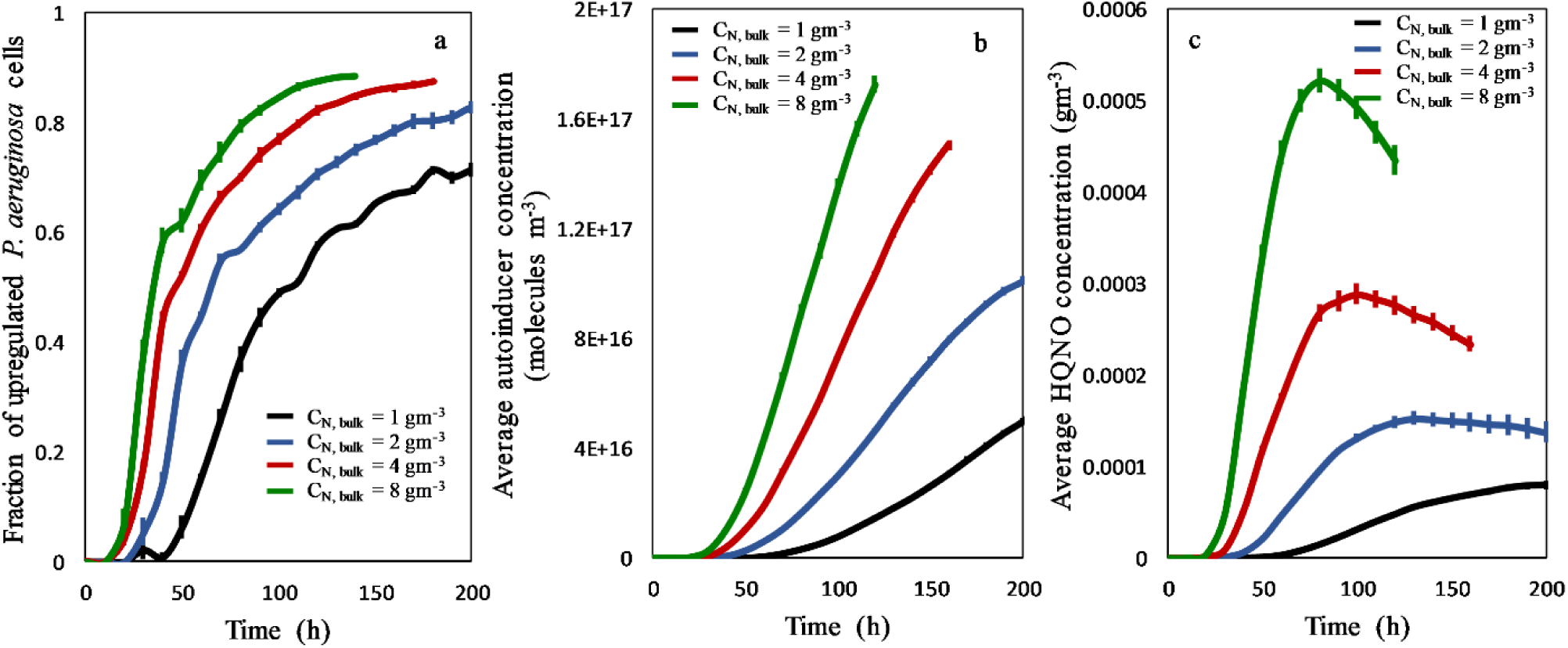
Quorum sensing and HQNO production. The fraction of upregulated *P. aeruginosa* cells (a), average autoinducer concentration (b), and average HQNO concentration (c) versus time for C_N,bulk_ = 8 gm^-3^ (green), 4 gm^-3^ (red), 2 gm^-3^ (blue), and 1 gm^-3^ (black). Data represent mean ± SEM of four separate simulations.

### 3.5 P. aeruginosa exoproducts induce phenotypic switching of wild-type S. aureus to SCVs

Wild-type *S. aureus* cells transform into small colony variants in the presence of HQNO released by *P. aeruginosa* cells in response to nutritional stress. The SCV biomass increased with time in the exponential phase, reached a peak, and then decreased (Fig. 5a). The rate of increase and the peak SCV biomass were higher for higher bulk nutrient concentrations, indicating that SCV biomass is directly related to the nutritional stress experienced by *P. aeruginosa* (Fig. 5a). Similar trends were observed for transformation events per wild-type *S. aureus* cell (Fig. 5b). Along these lines, HQNO available per *S. aureus* cell was higher for biofilms associated with higher bulk nutrient concentrations.

**Figure 5.**
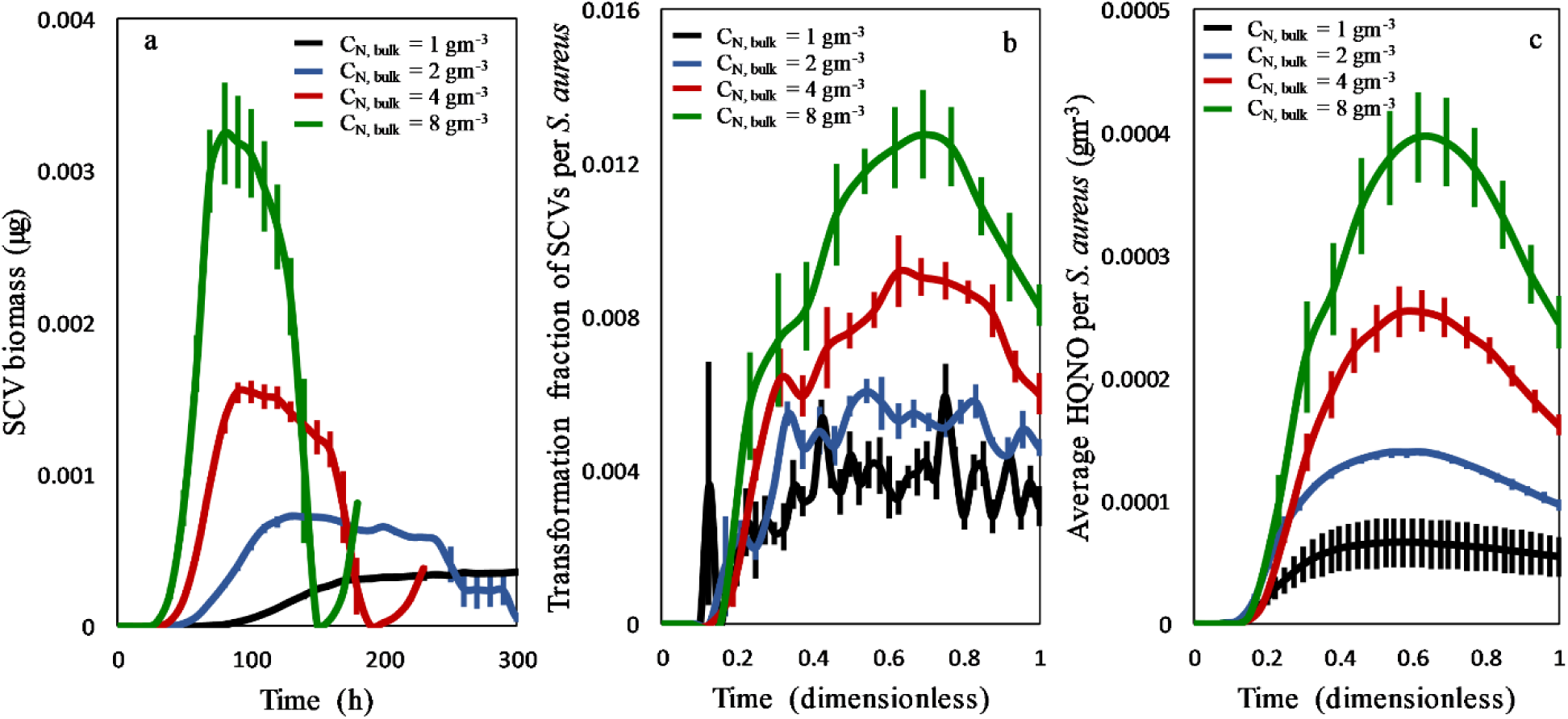
Growth dynamics of SCVs and HQNO-induced transformation of wild-type *S. aureus* into the SCV phenotype. Time evolution of SCV biomass (a), fraction of wild-type *S. aureus* transformed into SCVs (b), and average HQNO available per *S. aureus* cell (c) versus time for C_N,bulk_ = 8 gm^-3^ (green), 4 gm^-3^ (red), 2 gm^-3^ (blue), and 1 gm^-3^ (black). Data represent mean ± SEM of four separate simulations.

### 3.6 Synergistic effect of SCVs on P. aeruginosa

Taken altogether, our results indicate that nutritional stress for *P. aeruginosa* leads to enhanced secretion of HQNO, which in turn, potentiates phenotypic switching of wild-type *S. aureus* to dormant SCVs. We further propose that this phenotypic transformation has a synergistic influence, allowing *P. aeruginosa* to relieve nutritional stress. To test this hypothesis, we compared the relative *P. aeruginosa* biomasses in SCV^+^ and SCV^-^ biofilms. The ratio of relative *P aeruginosa* biomass in SCV^+^ to SCV^-^ biofilms was greater than one under all conditions tested (Fig. 6a), indicating that transformation of wild-type *S. aureus* into SCVs resulted in increased growth of *P. aeruginosa*. During the initial ∼70 h of growth, there’s virtually no excess *P. aeruginosa* biomass in the SCV^+^ biofilm, indicating that phenotypic switching is not effective up to this phase of growth. Subsequently, SCVs become effective as manifested by the monotonic increase in the ratio of relative *P. aeruginosa* biomass for the SCV^+^ to SCV^-^ biofilms. The excess *P. aeruginosa* biomass associated with the SCV^+^ biofilm is higher at higher bulk nutrient concentrations, indicating that SCVs are more effective in relieving nutritional stress for *P. aeruginosa* under these conditions. Combined with the previous observation that higher bulk nutrient concentrations are associated with higher nutritional stress (Fig. 3c), this indicates that phenotypic switching is upregulated in response to higher nutritional stress. Along these lines, nutrient availability to *P. aeruginosa* in the SCV^+^ biofilms was greater than that in the corresponding SCV^-^ biofilms with up to 40% excess nutrient available to *P. aeruginosa* in the SCV^+^ biofilm (Fig. 6b). This is a consequence of the fact that SCVs grow at a slower rate compared to wild-type *S. aureus*, and consequently consume lesser nutrient. Excess nutrient availability in the SCV^+^ biofilms is higher for higher bulk nutrient concentrations where *P. aeruginosa* is under greater nutritional stress. The overall dominance of *S. aureus* in the SCV^+^ biofilm decreased compared to that in the SCV^-^ biofilm, and the effect is more pronounced after ∼70 h of growth. Overall, these results suggest that *P. aeruginosa* uses HQNO-induced transformation of *S. aureus* into SCVs as a mechanism to relieve nutritional stress.

**Figure 6.**
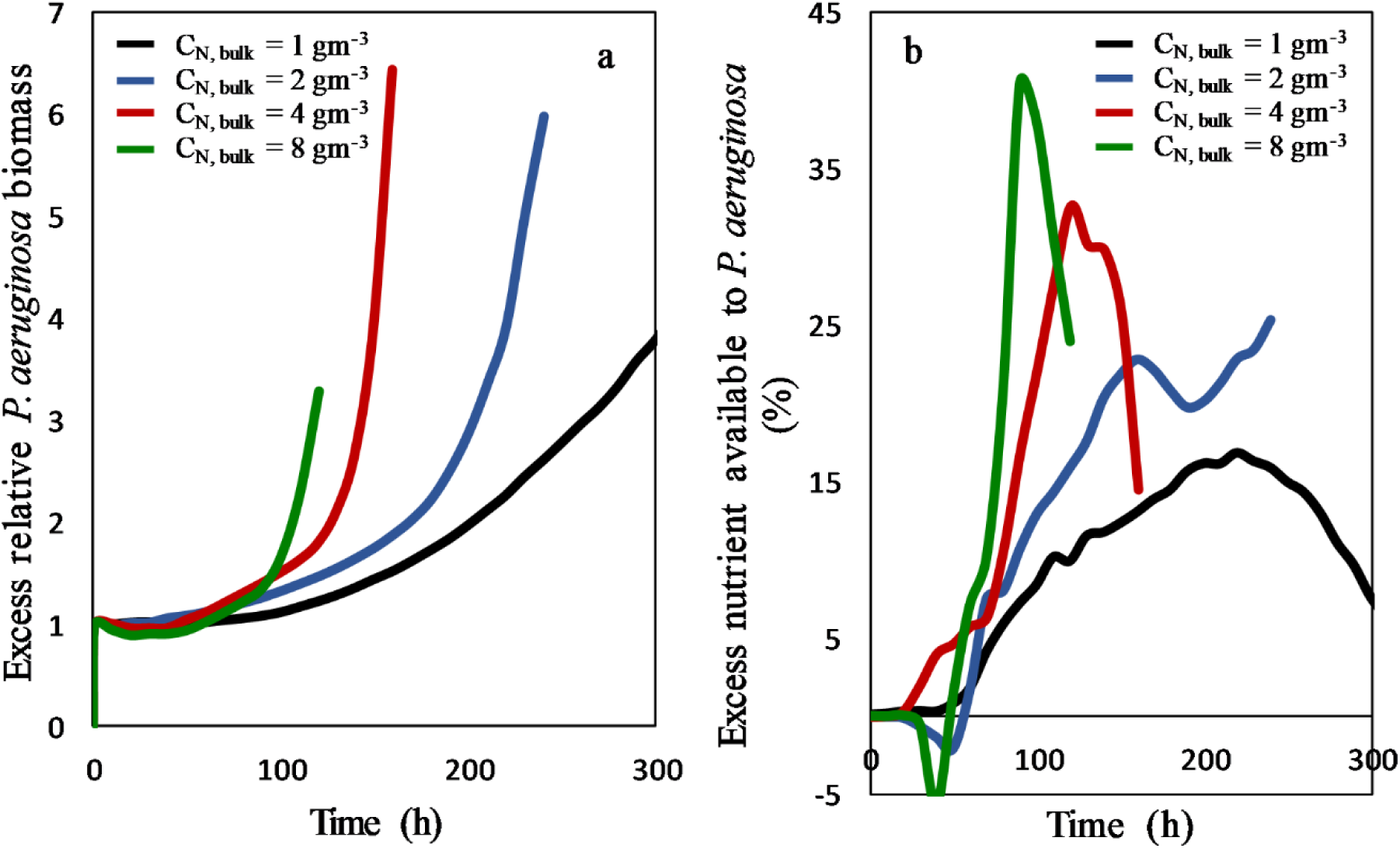
Synergistic effect of SCVs on *P. aeruginosa*. Ratio of relative *P. aeruginosa* biomasses for SCV^+^ and SCV^-^ biofilms (a), and % excess nutrient available to *P. aeruginosa* in SCV^+^ biofilms compared to SCV^-^ biofilms (b) versus time for C_N,bulk_ = 8 gm^-3^ (green), 4 gm^-3^ (red), 2 gm^-3^ (blue), and 1 gm^-3^ (black). Data represent mean ± SEM of four separate simulations.

## 4. Discussion

Polymicrobial biofilms exhibit unique characteristics typically not associated with monomicrobial biofilms, such as ecological competition for nutritional resources, interspecies QS, and phenotypic switching. This can lead to uneven distribution of nutrient availability, resulting in the eventual domination of one species over another as the biofilm matures. Using a 3D individual-based cellular automata model, we show that in a two-species biofilm comprised of *P. aeruginosa* and *S. aureus*, the former is under nutritional stress on account of its slower growth rates, and preferential localization towards the inner regions of the biofilm where nutrient availability is stunted due to diffusion limited transport. It has been shown in experimental systems that HQNO-induced phenotypic switching of wild-type *S. aureus* to the growth-impaired SCV phenotype allows *P. aeruginosa* to regain nutritional dominance. However, the mechanism that triggers release of HQNO is not fully understood. We hypothesize that unique biophysical characteristics associated with the heterogeneous architecture of the biofilm mode of growth may enhance the adaptive response of *P. aeruginosa* to nutritional stress. We propose that *P. aeruginosa* upregulates the release of HQNO as an adaptive stress response thereby inducing phenotypic switching in *S. aureus*. The resultant production of the metabolically challenged SCVs allows the nutritional stress for *P. aeruginosa* to be relieved. The release of HQNO and subsequent transformation of wild-type *S. aureus* to the quiescent SCV phenotype increases with increased nutritional stress. Once nutritional stress is relieved, HQNO secretion decreases. Based on these results, we propose that ecological competition in a two-species biofilm comprising *P. aeruginosa* and *S. aureus* proceeds via the following sequential phases:

- Phase 1: *Nutritional stress. P. aeruginosa* experiences nutritional stress as *S. aureus* dominates nutrient uptake. Nutritional stress increases with time as the biofilm matures, and the *S. aureus* subpopulation increases. Interestingly, greater nutritional stress is observed in biofilms associated with higher bulk nutrient concentrations.
- Phase 2: *Quorum sensing*. Maturation of the biofilm is associated with increased proportion of the *S. aureus* subpopulation resulting in increased autoinducer production. This results in a feedback-like mechanism in which upregulated cells secrete more autoinducer, causing a majority of the *P. aeruginosa* cells in the biofilm to switch to the upregulated state.
- Phase 3: *Secretion of exoproducts by P. aeruginosa*. In the mature biofilm, upregulated *P. aeruginosa* cells, under significant nutritional stress, release HQNO at an enhanced rate. HQNO concentrations within the biofilm are directly related to nutritional stress, indicative of a causal relationship.
- Phase 4: *Phenotypic switching*. Wild-type *S. aureus* cells undergo phenotypic switching to SCVs dependent on the local HQNO concentrations to which they are subjected. SCVs exhibit a markedly reduced nutrient uptake rate compared to wild-type *S. aureus* cells. Consequently, nutrient availability to *P. aeruginosa* increases.
- Phase 5: *Regulation*. As an increasing number of *S. aureus* cells transform into SCVs, autoinducer production diminishes, causing the QS mechanism to subside, resulting in lower transformation of *P. aeruginosa* cells from the downregulated to the upregulated states. This further reduces HQNO secretion, thereby lowering rates of phenotypic switching to SCVs.

Taken altogether, this work presents an effective cellular automata framework to investigate the influence of interspecies interactions on the growth dynamics of polymicrobial biofilms. Specific findings highlighted here could be further extended to investigate mechanisms of enhanced antibiotic resistance in multi-species bacterial biofilms.

## References

Abidi SH, Sherwani SK, Siddiqui TR, Bashir A, and Kazmi SU 2013 Drug resistance profile and biofilm forming potential of Pseudomonas aeruginosa isolated from contact lenses in Karachi-Pakistan. BMC Ophthalmol. 13

Abrudan MI, Smakman F, Grimbergen AJ, Westhoff S, Miller EL, Wezel GP Van, and Rozen DE 2015 Socially mediated induction and suppression of antibiosis during bacterial coexistence. Proc. Natl. Acad. Sci. U. S. A.

Armbruster CR et al. 2016 Staphylococcus aureus protein a mediates interspecies interactions at the cell surface of Pseudomonas aeruginosa. MBio 7

Armbruster CE, Hong W, Pang B, Weimer KED, Juneau RA, Turner J, and Edward Swords W 2010 Indirect pathogenicity of Haemophilus influenzae and Moraxella catarrhalis in Polymicrobial Otitis media occurs via interspecies quorum signaling. MBio 1

Azevedo AS, Almeida C, Melo LF, and Azevedo NF 2017 Impact of polymicrobial biofilms in catheter-associated urinary tract infections. Crit. Rev. Microbiol. 43 423–39

Bakkal S, Robinson SM, Ordonez CL, Waltz DA, and Riley MA 2010 Role of bacteriocins in mediating interactions of bacterial isolates taken from cystic fibrosis patients. Microbiology

Berbari EF, Osmon DR, Duffy MCT, Harmssen RNW, Mandrekar JN, Hanssen AD, and Steckelberg JM 2006 Outcome of Prosthetic Joint Infection in Patients with Rheumatoid Arthritis: The Impact of Medical and Surgical Therapy in 200 Episodes. Clin. Infect. Dis. 42 216–23

Bester E, Wolfaardt G, Joubert L, Garny K, and Saftic S 2005 Planktonic-Cell Yield of a *Pseudomonad* Biofilm. Appl. Environ. Microbiol. 71 7792–8

Chang I, Gilbert ES, Eliashberg N, and Keasling JD 2003 A three-dimensional, stochastic simulation of biofilm growth and transport-related factors that affect structure. Microbiology

Cohen TS et al. 2016 Staphylococcus aureus α toxin potentiates opportunistic bacterial lung infections. Sci. Transl. Med. 8 329ra31–329ra31

Déziel E, Lépine F, Milot S, He J, Mindrinos MN, Tompkins RG, and Rahme LG 2004 Analysis of Pseudomonas aeruginosa 4-hydroxy-2-alkylquinolines (HAQs) reveals a role for 4-hydroxy-2-heptylquinoline in cell-to-cell communication. Proc. Natl. Acad. Sci. U. S. A. 101 1339–44

Duus LM, Høiby N, Wang M, Schiøtz O, and Nørskov-Lauritsen N 2013 Bacteria of the genus Dyella can chronically colonise the airways of patients with cystic fibrosis and elicit a pronounced antibody response. Int. J. Med. Microbiol. 303 267–9

Emerenini BO, Hense BA, Kuttler C, and Eberl HJ 2015 A mathematical model of quorum sensing induced biofilm detachment. PLoS One

Eyoh AB edi. et al. 2014 Relationship between multiple drug resistance and biofilm formation in Staphylococcus aureus isolated from medical and non-medical personnel in Yaounde, Cameroon. Pan Afr. Med. J. 17 186

Fagerlind MG et al. 2012 Dynamic modelling of cell death during biofilm development. J. Theor. Biol. 295 23–36

Fazli M, Bjarnsholt T, Kirketerp-Møller K, Jørgensen B, Andersen AS, Krogfelt KA, Givskov M, and Tolker-Nielsen T 2009 Nonrandom distribution of Pseudomonas aeruginosa and Staphylococcus aureus in chronic wounds. J. Clin. Microbiol.

Filkins LM, Graber JA, Olson DG, Dolben EL, Lynd LR, Bhuju S, and O’Toole GA 2015 Coculture of Staphylococcus aureus with Pseudomonas aeruginosa drives S. aureus towards fermentative metabolism and reduced viability in a cystic fibrosis model. J. Bacteriol. 197 2252–64

Filkins LM and O’Toole GA 2015 Cystic Fibrosis Lung Infections: Polymicrobial, Complex, and Hard to Treat. PLOS Pathog. 11 e1005258

Fozard JA, Lees M, King JR, and Logan BS 2012 Inhibition of quorum sensing in a computational biofilm simulation. Biosystems 109 105–14

Gardner SE and Frantz RA 2008 Wound bioburden and infection-related complications in diabetic foot ulcers. Biol. Res. Nurs. 10 44–53

Gjødsbøl K, Christensen JJ, Karlsmark T, Jørgensen B, Klein BM, and Krogfelt KA 2006 Multiple bacterial species reside in chronic wounds: A longitudinal study. Int. Wound J. 3 225–31

Guélon T, Mathias JD, and Deffuant G 2012 Influence of spatial structure on effective nutrient diffusion in bacterial biofilms. J. Biol. Phys. 38 573–88

Gutierrez Jauregui R, Fleige H, Bubke A, Rohde M, Weiss S, and Förster R 2019 IL-1β Promotes Staphylococcus aureus Biofilms on Implants in vivo. Front. Immunol. 10 1082

Hall-Stoodley L, Costerton JW, and Stoodley P 2004 Bacterial biofilms: From the natural environment to infectious diseases. Nat. Rev. Microbiol.

Haruta S, Kato S, Yamamoto K, and Igarashi Y 2009 Intertwined interspecies relationships: Approaches to untangle the microbial network. Environ. Microbiol.

Hoffman LR et al. 2006 Selection for Staphylococcus aureus small-colony variants due to growth in the presence of Pseudomonas aeruginosa. Proc. Natl. Acad. Sci. U. S. A.

Hoffman LR et al. 2010 Nutrient Availability as a Mechanism for Selection of Antibiotic Tolerant Pseudomonas aeruginosa within the CF Airway. PLoS Pathog. 6 e1000712

Høiby N et al. 2017 Diagnosis of biofilm infections in cystic fibrosis patients. APMIS 125 339–43

Hunt SM, Werner EM, Huang B, Hamilton MA, and Stewart PS 2004 Hypothesis for the role of nutrient starvation in biofilm detachment. Appl. Environ. Microbiol.

Ito A, Taniuchi A, May T, Kawata K, and Okabe S 2009 Increased antibiotic resistance of Escherichia coli in mature biofilms. Appl. Environ. Microbiol. 75 4093–100

Korgaonkar A, Trivedi U, Rumbaugh KP, and Whiteley M 2013 Community surveillance enhances Pseudomonas aeruginosa virulence during polymicrobial infection. Proc. Natl. Acad. Sci. U. S. A. 110 1059–64

Korgaonkar AK and Whiteley M 2011 Pseudomonas aeruginosa enhances production of an antimicrobial in response to N-acetylglucosamine and peptidoglycan. J. Bacteriol. 193 909–17

Kreft J-U, Booth G, and Wimpenny JWT 1998 BacSim, a simulator for individual-based modelling of bacterial colony growth. Microbiology 144 3275–87

Kroukamp O, Dumitrache RG, and Wolfaardt GM 2010 Pronounced Effect of the Nature of the Inoculum on Biofilm Development in Flow Systems. Appl. Environ. Microbiol. 76 6025–31

Langebrake JB, Dilanji GE, Hagen SJ, and Leenheer P De 2014 Traveling waves in response to a diffusing quorum sensing signal in spatially-extended bacterial colonies. J. Theor. Biol. 363 53–61

Lau GW, Hassett DJ, Ran H, and Kong F 2004 The role of pyocyanin in Pseudomonas aeruginosa infection. Trends Mol. Med.

Lightbown JW and Jackson FL 1956 Inhibition of cytochrome systems of heart muscle and certain bacteria by the antagonists of dihydrostreptomycin: 2-alkyl-4-hydroxyquinoline N-oxides. Biochem. J. 63 130–7

Machineni L, Rajapantul A, Nandamuri V, and Pawar PD 2017 Influence of Nutrient Availability and Quorum Sensing on the Formation of Metabolically Inactive Microcolonies Within Structurally Heterogeneous Bacterial Biofilms: An Individual-Based 3D Cellular Automata Model. Bull. Math. Biol.

Machineni L, Reddy CT, Nandamuri V, and Pawar PD 2018 A 3D individual-based model to investigate the spatially heterogeneous response of bacterial biofilms to antimicrobial agents.; in: Mathematical Methods in the Applied Sciences

Martin B, Tamanai-Shacoori Z, Bronsard J, Ginguené F, Meuric V, Mahé F, and Bonnaure-Mallet M 2017 A new mathematical model of bacterial interactions in two-species oral biofilms. PLoS One 12 e0173153

Mazumdar V, Amar S, and Segrè D 2013 Metabolic Proximity in the Order of Colonization of a Microbial Community. PLoS One 8 e77617

McBirney SE, Trinh K, Wong-Beringer A, and Armani AM 2016 Wavelength-normalized spectroscopic analysis of Staphylococcus aureus and Pseudomonas aeruginosa growth rates. Biomed. Opt. Express

McDaniel MS, Schoeb T, and Swords WE 2020 Cooperativity between Stenotrophomonas maltophilia and Pseudomonas aeruginosa during Polymicrobial Airway Infections. Infect. Immun. 88

Mitchell G, Séguin DL, Asselin AE, Déziel E, Cantin AM, Frost EH, Michaud S, and Malouin F 2010 Staphylococcus aureus sigma B-dependent emergence of small-colony variants and biofilm production following exposure to Pseudomonas aeruginosa 4-hydroxy-2-heptylquinoline-N-oxide. BMC Microbiol.

Moghadam SO, Pourmand MR, and Aminharati F 2014 Biofilm formation and antimicrobial resistance in methicillin-resistant staphylococcus aureus isolated from burn patients, Iran. J. Infect. Dev. Ctries. 8 1511–7

Nadell CD, Drescher K, and Foster KR 2016 Spatial structure, cooperation and competition in biofilms. Nat. Rev. Microbiol.

Nguyen AT, Jones JW, Ruge MA, Kane MA, and Oglesby-Sherrouse AG 2015 Iron depletion enhances production of antimicrobials by Pseudomonas aeruginosa. J. Bacteriol.

Nyström T 2001 Not quite dead enough: On bacterial life, culturability, senescence, and death. Arch. Microbiol.

Nyström T 2003 Conditional senescence in bacteria: Death of the immortals. Mol. Microbiol.

Oliveira NM, Martinez-Garcia E, Xavier J, Durham WM, Kolter R, Kim W, and Foster KR 2015 Biofilm formation as a response to ecological competition. PLoS Biol.

Parijs I and Steenackers HP 2018 Competitive inter-species interactions underlie the increased antimicrobial tolerance in multispecies brewery biofilms. ISME J.

Perez AC et al. 2014 Residence of *Streptococcus pneumoniae* and *Moraxella catarrhalis* within polymicrobial biofilm promotes antibiotic resistance and bacterial persistence *in vivo*. Pathog. Dis. 70 280–8

Petroff AP, Wu T Di, Liang B, Mui J, Guerquin-Kern JL, Vali H, Rothman DH, and Bosak T 2011 Reaction-diffusion model of nutrient uptake in a biofilm: Theory and experiment. J. Theor. Biol. 289 90–5

Phalak P, Chen J, Carlson RP, and Henson MA 2016 Metabolic modeling of a chronic wound biofilm consortium predicts spatial partitioning of bacterial species. BMC Syst. Biol.

Picioreanu C, Kreft JU, and Loosdrecht MCM Van 2004 Particle-based multidimensional multispecies biofilm model. Appl. Environ. Microbiol.

Picioreanu C, Loosdrecht MCM Van, and Heijnen JJ 1998 A new combined differential-discrete cellular automaton approach for biofilm modeling: Application for growth in gel beads. Biotechnol. Bioeng.

Proctor RA, Eiff C von, Kahl BC, Becker K, McNamara P, Herrmann M, and Peters G 2006 Small colony variants: A pathogenic form of bacteria that facilitates persistent and recurrent infections. Nat. Rev. Microbiol.

Pulimood S, Ganesan L, Alangaden G, and Chandrasekar P 2002 Polymicrobial candidemia. Diagn. Microbiol. Infect. Dis.

Qi L et al. 2016 Relationship between Antibiotic Resistance, Biofilm Formation, and Biofilm-Specific Resistance in Acinetobacter baumannii. Front. Microbiol. 7 483

Sena NT, Gomes BPFA, Vianna ME, Berber VB, Zaia AA, Ferraz CCR, and Souza-Filho FJ 2006 In vitro antimicrobial activity of sodium hypochlorite and chlorhexidine against selected single-species biofilms. Int. Endod. J. 39 878–85

Stewart PS 1998 A review of experimental measurements of effective diffusive permeabilities and effective diffusion coefficients in biofilms. Biotechnol. Bioeng. 59 261–72

Stewart PS 2003 Diffusion in biofilms. J. Bacteriol. 185 1485–91

Swidsinski A, Verstraelen H, Loening-Baucke V, Swidsinski S, Mendling W, and Halwani Z 2013 Presence of a Polymicrobial Endometrial Biofilm in Patients with Bacterial Vaginosis. PLoS One 8

Tilman D 1977 Resource Competition between Plankton Algae: An Experimental and Theoretical Approach. Ecology 58 338–48

Traxler MF, Watrous JD, Alexandrov T, Dorrestein PC, and Kolter R 2013 Interspecies interactions stimulate diversification of the Streptomyces coelicolor secreted metabolome. MBio

Vuotto C, Longo F, Balice M, Donelli G, and Varaldo P 2014 Antibiotic Resistance Related to Biofilm Formation in Klebsiella pneumoniae. Pathogens 3 743–58

Wijesinghe G, Dilhari A, Gayani B, Kottegoda N, Samaranayake L, and Weerasekera M 2019 Influence of Laboratory Culture Media on in vitro Growth, Adhesion, and Biofilm Formation of Pseudomonas aeruginosa and Staphylococcus aureus. Med. Princ. Pract.

Woods J, Boegli L, Kirker KR, Agostinho AM, Durch AM, deLancey Pulcini E, Stewart PS, and James GA 2012 Development and application of a polymicrobial, in vitro, wound biofilm model. J. Appl. Microbiol. 112 998–1006

